# Effect of *Sargassum* on the Behavior and Survival of the Earthworm *Eisenia fetida*

**DOI:** 10.1101/2023.09.08.556937

**Authors:** Mayela Martínez-Cano, Ana E. Dorantes-Acosta, Rogelio Lara-González, Enrique Salgado-Hernández, Angel I. Ortiz-Ceballos

**Affiliations:** Instituto de Biotecnología y Ecología Aplicada (INBIOTECA), Universidad Veracruzana. Av. de las Culturas Veracruzanas 101. Col. E. Zapata. CP 91090, Xalapa, Veracruz, México

**Keywords:** seaweed, heavy metals, polyphenols, biofertilizers, earthworm compost, oligochaetes, biosolids, pollution

## Abstract

In the past decade, *Sargassum natans* and *S. fluitans* have massively reached the Mexican Caribbean shores. As a result, *Sargassum* confinement sites may be affecting the soil quality and health in coastal ecosystems and agroecosystems. The impact of *Sargassum* (e.g., polyphenols) on soil biodiversity has not yet been evaluated. Terrestrial ecotoxicological tests use the epigean earthworm *Eisenia fetida* as a model organism to assess the function of soil habitats. This study evaluated the behavior and survival of the earthworm *E. fetida* exposed to five *Sargassum* concentrations (0, 25, 50, 75, and 100 %) using two toxicological tests. The avoidance test showed that *E. fetida* repelled (>80 %) a diet with 100 % *Sargassum*. In contrast, the acute test recorded a low mortality; however, the growth of *E. fetida* was lower with increasing *Sargassum* concentrations. It is concluded that the ability to repel and *E. fetida* biomass are early warning bioindicators to predict the environmental risk of *Sargassum* in soil. Therefore, it is relevant to determine the potential risk of using earthworm compost and *Sargassum* leachates as biofertilizers in agroecosystems.

## Introduction

Seaweed comprises a set of marine macroalgae of the genus *Sargassum*, including *S. natans* and *S. fluitans*, (Gower et al. 2013). and is the natural habitat of various marine species (Suarez-Castillo, 2008; Baker et al. 2018). However, human activity has led to changes in the chemical characteristics of the Atlantic Ocean; particularly, since 2011, the coasts of the Mexican Caribbean Sea started recording unusually high amounts of *Sargassum*. (Wang et al. 2019). Algae blooms produce eutrophication, reflected as higher temperature and available nutrient levels, and low mobility and dissolved oxygen (Rodríguez-Martínez et al. 2019). Sargassum landings are on the rise and have become an issue due to the mortality of fauna associated with *Sargassum* (Paredes Rangel, 2020). In addition to environmental problems, *Sargassum* has affected human health and tourism (Muñoz, 2019) at local and regional levels (Resiere et al. 2019). For this reason, several strategies have been implemented to collect the large volumes of *Sargassum* biomass reaching beaches and dispose of it in landfills in natural ecosystems (forests, mangroves, beaches), agroecosystems, and suburban areas. It has been suggested that *Sargassum* be used as a biofertilizer (Khan et al. 2009; Nosiri et al. 2019), animal feed (Choi et al. 2020; Gojon et al. 1998), and in biogas production (Salgado-Hernández et al. 2023; Thompson et al. 2021), for cosmetics (Nurjanah et al., 2018), pharmaceuticals, construction, and textile production (Oyesiku and Egunyomi 2014), among others. However, it has been documented that the *Sargassum* biomass from the Mexican Caribbean contains heavy metals (Rodríguez Martínez et al. 2020) and polyphenols (Salgado-Hernández et al. 2023). This poses a risk to soil biodiversity (Ekschmitt et al. 2003), aquifers, and human health in areas where *Sargassum* is landfilled (SEMARNAT 2021).

Soil ecotoxicity studies have been widely used to determine the effect of pollution by heavy metals, pesticides, hydrocarbons, herbicides, and microplastics (Zhang et al. 2014; Rodríguez-Seijo 2018; Li et al. 2020). There are currently well-established indicator organisms, including earthworms, for short- and long-term ecological toxicity monitoring purposes(Yeardley et al. 1996; Graefe and Tischer 2011; Castracani et al. 2015). Earthworms are clitellate oligochaetes known as “ecosystem engineers” for improving the physical, chemical, and biological properties of soil and boosting plant productivity (Bohlen et al. 2004; Domínguez and Aira 2009); that is, they represent soil biocenosis in ecotoxicological studies (Paoletti 1999; ISO 2008). Standardized ecotoxicology methods use the epigean earthworm *Eisenia fetida* as a model organism to record repellency and mortality (OECD 1984; ISO 2008). The impact of *Sargassum* on soil biodiversity has yet to be studied. Thus, the objective of this study was to evaluate experimentally the sensitivity of the tropical epigean earthworm *Eisenia fetida* exposed to different *Sargassum* concentrations.

## Materials and Methods

### Sargassum Collection and Processing

*Sargassum* was collected in Playa del Carmen (20º37’19.99” N, 87º4’9.98” W), Quintana Roo, Mexico. The foliage of other marine species was removed, and then *Sargassum* was placed in a cooler. In the laboratory, it was kept at -5 ºC; before the experiments (6 hrs), it was thawed until it reached an ambient temperature of 25 ºC The collected *Sargassum* was characterized by containing (%) 1.68 N, 1.47 K, 9.98 Ca, 3.36 Na, 3.11 H, and 24.7 C, in addition to (mg/kg, dry weight) 849.19 P, 0.74 Mg, 6037.65 S, 5.08 Cu, 19.04 Mn, 175.90 Fe, and 30.76 Zn.

### Coffee Pulp

Since several years ago, coffee pulp from the coffee regions of Mexico and Latin America has been used as an organic substratum for breeding *E. fetida* (Dávila and Ramírez 1996). The disgestion product of coffee pulp ingested by *E. fetida* is the “earthworm compost”, a biofertilizer for agricultural use (NMX-FF-109-SCFI-2008).

The nutritional composition of coffee pulp contains (mg/kg) 0.004 to 17 N, 0.01 to 2.48 P, and 0.016 to 25.13 K, along with a 31.43 C/N ratio (Aranda et al. 2000; Braham and Bressani 1978; Fierro and Morales-Ramos 2018; Noriega-Salazar et al. 2008; Puerta Quintero and Arias, 2011; Serna-Jimenez et al. 2018).

We collected 60 kg of coffee pulp from the company “Lombricomposta Olmos” for use as substrate (habitat and food). In the greenhouse, coffee pulp was kept in plastic boxes (40 cm x 50 cm x 10 cm) at 80 % humidity and room temperature until use in the ecotoxicology experiments.

### *Eisenia fetida* Earthworms

The company “Lombricomposta Olmos” (San Marcos, Veracruz, Mexico) provided 300 *E. fetida* juvenile earthworms used in the present study. Earthworms used in the experiment were maintained in a greenhouse in plastic boxes (40 cm x 50 cm x 10 cm) containing coffee pulp at 80 % humidity and 24 °C.

### Avoidance Test Experimental Setup

The laboratory experiment was conducted in the Institute of Biotechnology and Applied Ecology at Universidad Veracruzana, Mexico, based on the ISO 1751-2 standard (ISO 2008). Using a completely randomized design, we evaluated five *Sargassum* concentrations or treatments (0, 25, 50, 75, and 100 %), with five replicates each, using glass containers (20 cm x 10 cm x 10 cm) as replicates. The glass containers were divided into two sections with a plastic separator (10 cm x 10 cm): A (control = coffee pulp) and B (test = *Sargassum* + coffee pulp). In each control section (A) of the five treatments, coffee pulp (200 g) was placed up to 5 cm high. In the test section (B), a homogeneous mixture of *Sargassum* and coffee pulp corresponding to each treatment was placed at the following *Sargassum*:coffee pulp ratios: 0:0, 25:75, 50:50, 75:25, and 100:0. Afterward, the plastic separator was removed from all containers; then, 10 *Eisenia fetida* individuals were introduced in the gap between sections A and B; earthworms weighed 2000 mg ± 3 mg on average; then, the containers were covered to maintain humidity and temperature and prevent earthworms from escaping.

After 48 hours of culture (end of the experiment), the separator between sections A and B was reinstalled in all replicas or glass containers; then, the earthworms in each section were manually collected, washed, dried, weighed, and counted.

The relative effect of *Sargassum* concentration (%) was calculated by comparing the average number of earthworms in the test substrate (*Sargassum*) with the number in the control (coffee pulp) [0 % avoidance was considered a negative response (= earthworms prefer the test substrate)]. The following equation was used (ISO 2008):

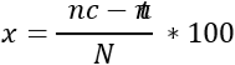

Where *X* = evasion expressed as a percentage; *nc* = number of worms in the control substrate; *nt* = number of worms in the test substrate; *N* = total number of worms in the test. The final *E. fetida* biomass data was evaluated using an analysis of variance (ANOVA), while the differences between means were analyzed with Tukey’s HSD test.

### Acute Toxicity Test

The acute test was carried out In the laboratory for 14 days based on a completely randomized design as per the ISO 11268-1 standard (ISO 2012). Five treatments were studied: 0 % = 100 % coffee pulp + 0 % *Sargassum*: 25 % = 75 % coffee pulp + 25 % *Sargassum*; 50 % = 50 % coffee pulp + 50 % *Sargassum*; 75 % = 25 % coffee pulp + 75 % *Sargassum*; and 100 % = 0 % coffee pulp + 100 % *Sargassum*. Each treatment included five replicas or glass containers (10 cm x 20 cm x 10 cm) with 10 *E. fetida* earthworms per replicate.

In the glass containers, according to the respective treatment, 200 g of a homogeneous mixture of *Sargassum* and coffee pulp was placed; then, 10 juvenile earthworms of similar biomass (2000.0 mg ± 5 mg) were introduced; afterward, the containers were covered with a plastic film to preserve temperature and humidity and prevent earthworms from escaping. The containers were kept for 14 days at room temperature and constant humidity (90 %).

On days 7 and 14 of the experiment, the living earthworms were collected, washed, dried, weighed, and counted. Earthworms that did not respond to a mechanical stimulus were considered dead.

### Data Analysis

Biomass data were evaluated using a one-way analysis of variance (ANOVA). Differences between means were analyzed with Tukey’s HSD test. The statistical analysis was carried out using the STATGRAPHICS Centurion software.

## Results

In the avoidance test, all earthworms (*n* = 250) were alive at 48 hrs in the five *Sargassum* treatments. Fresh *E. fetida* biomass varied significantly between treatments (*F* = 4.68, *p* = 0.004). Figure 1 shows that earthworms recorded a lower biomass in the control (0 % *Sargassum*) than in the *Sargassum* treatments (25 %, 50 %, 75 %, and 100 %;); furthermores, earthworms showed no changes in behavior or signs of external damage.

**Figure 1.**
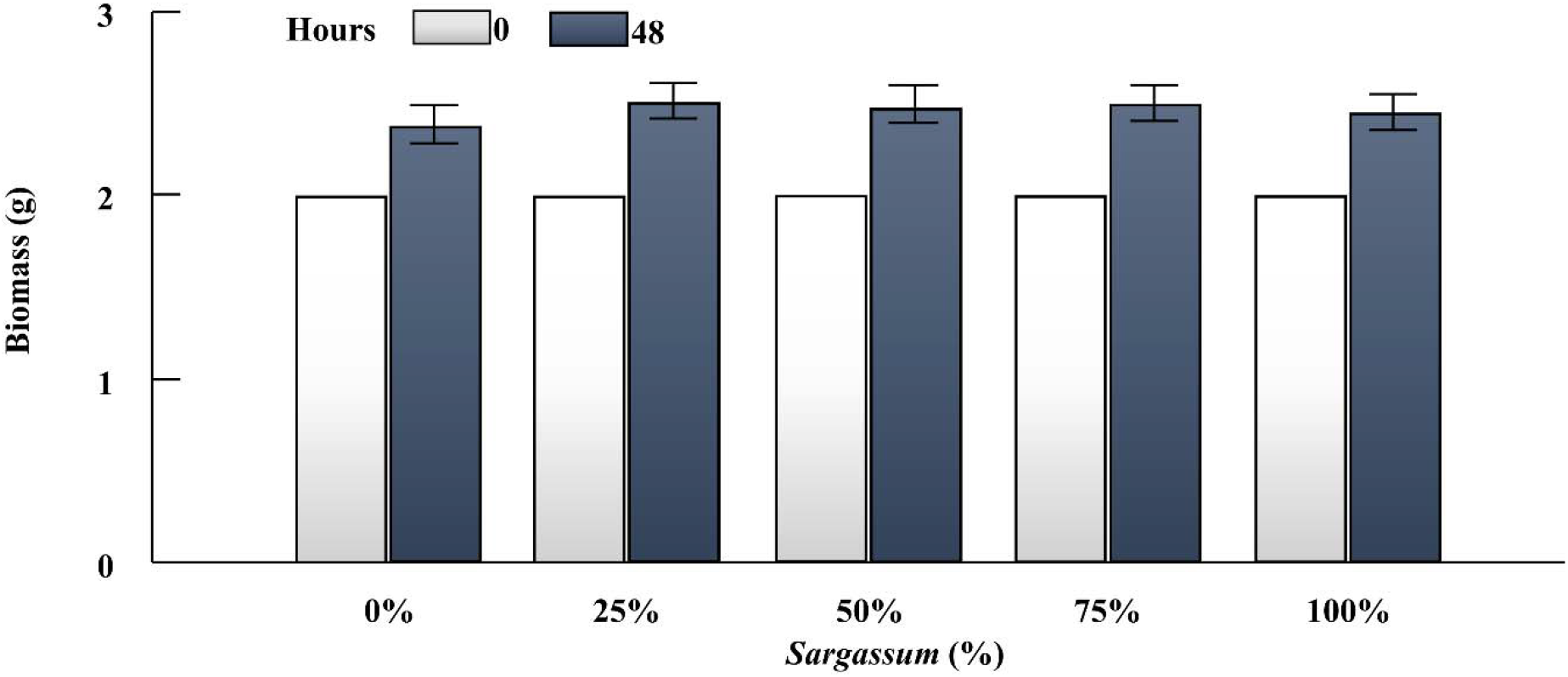
Initial and final average biomass of the tropical epigean earthworm *Eisenia fetida* after 48 hrs of exposure to five *Sargassum* diets. Values are averages; different letters indicate significant differences at *p* < 0.05 (Tukey’s HSD test); vertical lines represent the standard error (*n* = 5).

However, the results showed a significant variation in the behavior of *E. fetida* between *Sargassum* treatments (*F* = 5.7, *p* = 0.003). Earthworms repelled the 100 % Sargassum treatment as habitat (>80 %) but did not avoid the 25 %, 50 %, and 75 % *Sargassum* treatment (Figure 2).

**Figure 2.**
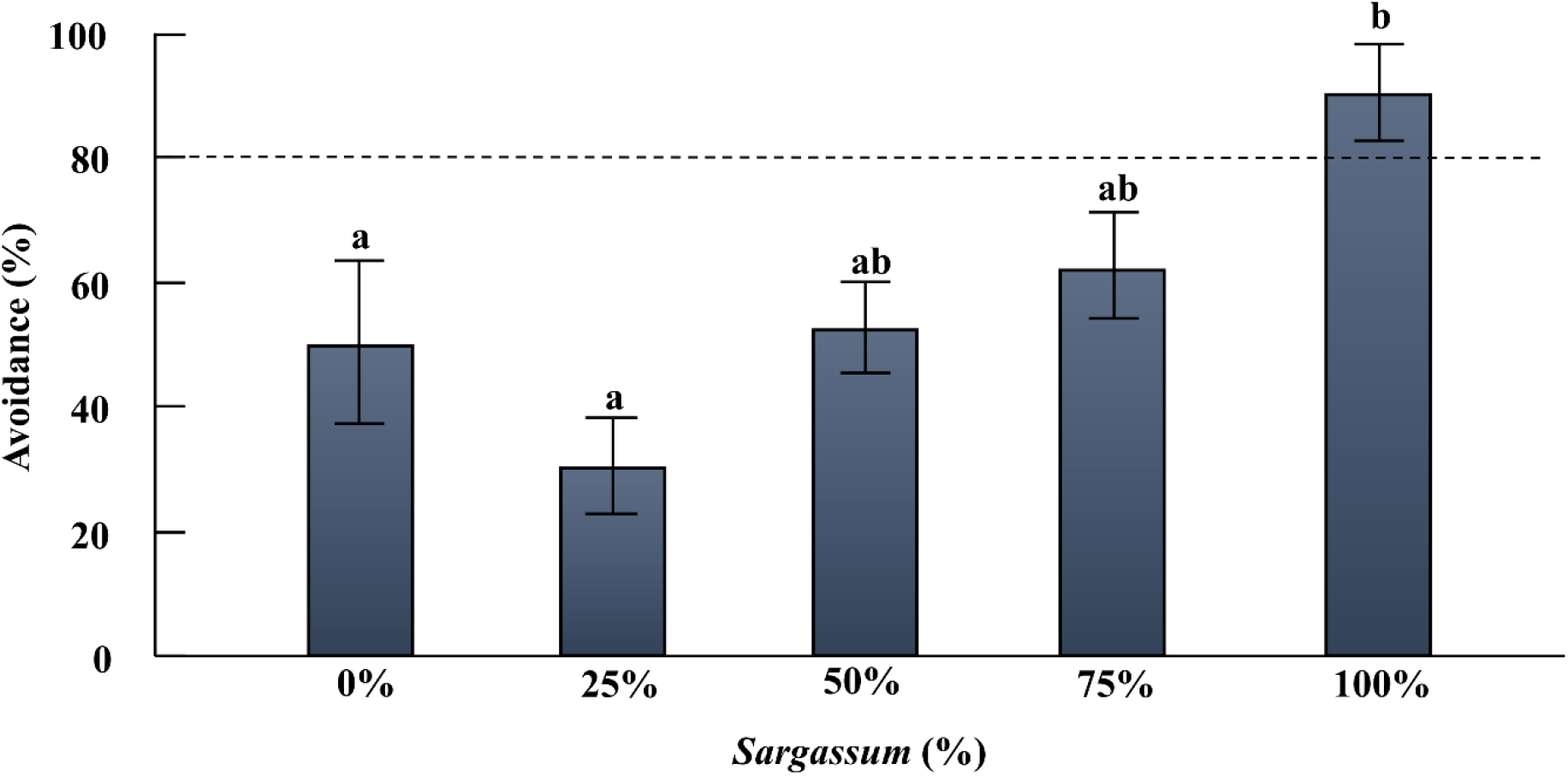
Avoidance response (%) of the tropical epigean earthworm *Eisenia fetida* after 48 hrs of exposure to five *Sargassum* diets. Values are average; vertical lines represent the standard error (*n* = 5). Different letters represent significant differences at *p* < 0.05 (Tukey’s HSD test).

In the acute test, *E. fetida* mortality at day 14 was similar between treatments; i.e., the 0 %, 75 %, and 100 % *Sargassum* treatments produced no deaths, but in the 25 % and 50 % treatments, 2 and 1 earthworms were found dead, respectively (Figure 3).

**Figure 3.**
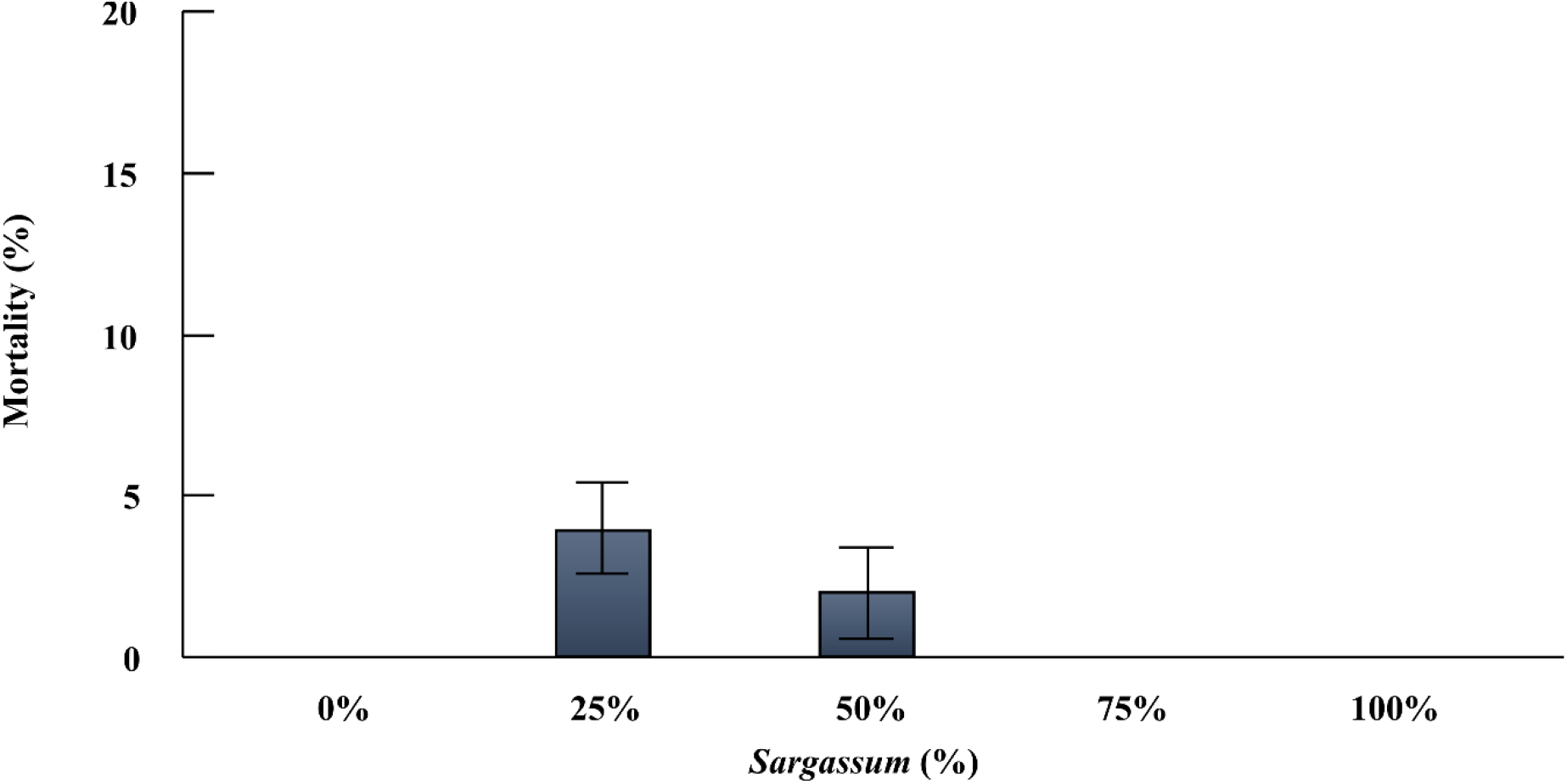
Average mortality of the tropical epigean earthworm *Eisenia fetida* at days 7 and 14 of exposure to five *Sargassum* diets. Values are averages and vertical lines represent the standard error (*n* = 5).

The biomass of *E. fetida* at days 7 and 14 varied significantly between treatments (*F* = 8.03, *p* = 0.0005 and *F* = 5.40, *p* = 0.004, respectively). At days 7 and 14, *E. fetida* biomass was higher in the 25 % *Sargassum* treatment and lower in the 100 % treatment (Figure 4).

**Figure 4.**
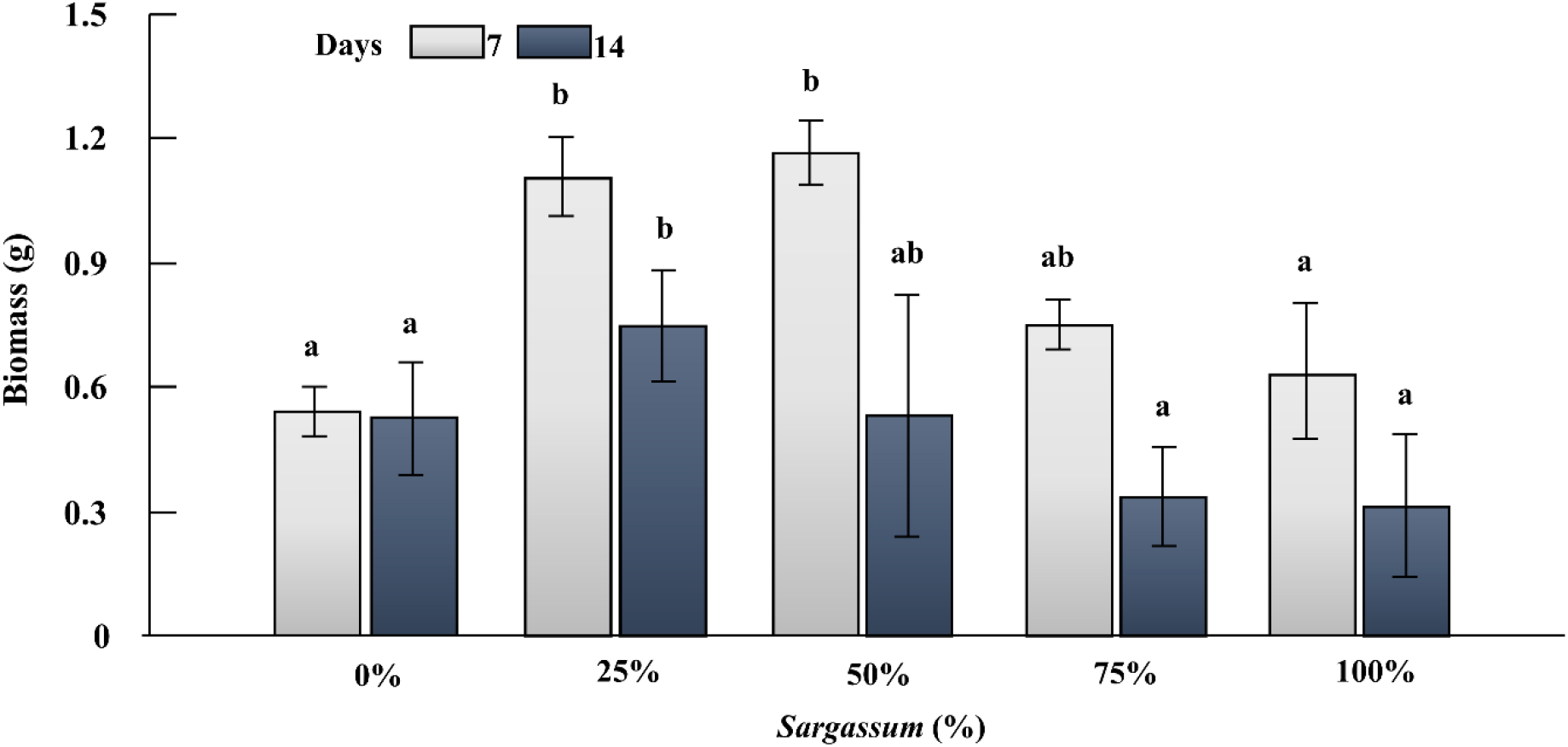
Initial and final average biomass of the tropical epigean earthworm *Eisenia fetida* after 7 days of exposure to five *Sargassum* diets. Vertical lines represent the standard error (*n* = 5). Different letters represent significant differences at *p* < 0.05 (Tukey’s HSD test).

## Discussion

Terrestrial ecotoxicological test systems are used to assess the effects of pollutants on soil (ISO 2008) and predict their effect on agroecosystems and ecosystems (Aulakh et al. 2022). In the present study, the results of the avoidance and mortality trial showed a significant sensitivity to *Sargassum* exposure over a short period.

*Sargassum* collected in the Mexican Caribbean has tested positive for heavy metals (Oyesiku and Egunyomi, 2014; Saldarriaga Hernandez et al. 2021; Tonon et al. 2022). *E. fetida* has also been reported to repel (>80 %) sodium (>2000 mg/kg NaCl; Owojori et al. 2009), Zn (400 mg/kg; Gao et al. 2016), and Cu (0.5 mg/kg; Demuynck et al. 2016) This may explain the results of the present study by demonstrating that the tropical epigean earthworm *E. fetida* rejected the 100 % *Sargassum* treatment.

The acute test results showed a non-significant mortality of *E. fetida*, suggesting that *Sargassum* did not produce a lethal effect, similar to the findings recorded in other studies with As, Fe, Al, Cr, Mn, Zn, Cu, and Na (Dzul Caamal et al. 2020; Lee et al. 2013; Owojori et al. 2009). It is known that enzymatic activity and antioxidant defense protect earthworms from oxidative stress associated with heavy metals, herbicides, and other chemicals (Owagboriaye et al. 2019). The low mortality of *E. fetida* is probably due to the heavy metals accumulated in *Sargassum* being below the average lethal concentration recorded with As, Cr, Pb, Cd, Ni, Zn, and Cu in natural and artificial polluted soils (Aulakh et al. 2022); furthermore, immobility, stiffness, or pigmentation changes were not observed (Piearce et al. 2002; Miglani and Bisht 2019; Chauhan and Suthar 2020). In this regard, it has been suggested that the accumulation-excretion patterns of heavy metals are highly variable across earthworm species. These factors are also influenced by the environment, including exposure time, soil type, organic matter, and others (Tözsér et al. 2022). Therefore, our results suggest evaluating prolonged exposure to *Sargassum* to determine whether it produces a delayed effect on *E. fetida* (Guimarães et al. 2023).

The avoidance and mortality test recorded a significant change in the biomass of *E. fetida* associated with the *Sargassum* diet. These results can be explained because *Sargassum* has a higher nutritional quality than coffee pulp. However, by increasing the *Sargassum* content in the diet, the growth of *E. fetida* decreased, maybe due to the presence of heavy metals (Roidríguez Martínez et al. 2020; Tonon et al. 2022) and polyphenols (Salgado Hernandez et al. 2023). Phenols are inhibitors of microorganisms, which have a symbiotic relationship with earthworms (Edwards and Fletcher 1988). However, phenols do not have a lethal effect, as earthworms produce active metabolites called “drilodefensins” to offset the inhibitory effect of polyphenols (Liebeke et al. 2015). It has also been suggested that the symbiotic bacteria-phage interaction (more bacteria, less viruses) in the gut of earthworms facilitates the adaptation of microbiota to stress by pollutants (Kirsch et al. 2021; Knowles et al. 2016; Xia et al. 2023). Phages found as virions within microbial cells provide resistance to pollutants present at low concentrations (Xia et al. 2023); for example, a cooperative bacteria-phage relationship has been reported associated with low benzene(a)pyrene (BaP) levels (Xia et al. 2023). The above likely explains why *E. fetida* fed a diet with low (25 %) to intermediate (50 %) *Sargassum* content (i.e., low levels of polyphenols and heavy metals) increased its biomass by 93 % and 85 %, respectively; in contrast, a diet with high (75 %) and very high (100 %) *Sargassum* content produced a lower increase in the *E. fetida* biomass (55 % and 48 %, respectively), similar to the weight gain observed with coffee pulp alone (55 %). These results point to the need to evaluate the effects of prolonged exposure to *Sargassum* on *E. fetida* using the changes in the antioxidant system and enzymatic activity as indicators (Hoffman 2003; Kaur et al. 2022).

Finally, it has been suggested to use the earthworms *E. fetida, E. andrei, Eudrilus eugeniae*, and *Perionix excavations* to neutralize the effects of toxic substances in polluted sites (Gudeta et al. 2023; Sharma and Garg 2018). Then, a potentially relevant management protocol (in *Sargassum* landfills) would be inoculating and propagating *E. fetida* and incorporating coffee pulp or other organic materials (agricultural, urban, and industrial) to accumulate and biotransform heavy metals, polyphenols, and other pollutants derived from *Sargassum* (Aulakh et al. 2022; Gudeta et al. 2023; Sharma & Garg 2018). However, the ecological relevance of *E. fetida* for evaluating soil quality has been questioned, as it only thrives in organic waste (biosolids) and does not inhabit the soil; instead, geophagous (endogean) species have been suggested as ideal and most susceptible to soil pollutants (Aulakh et al. 2022). Therefore, it is relevant to use native endogean earthworms from the Yucatan Peninsula, such as *Balanteodrilus pearsei, Diplotrema oxcutzcabensis*, and *Mayadrilus calakmulensis* (Fragoso et al. 2016) to evaluate the soil condition of ecosystems and agroecosystems in areas where *Sargassum* collected on the Mexican Caribbean coasts is deposited.

## Conclusions

The results of the present work showed that *E. fetida* significantly repelled the *Sargassum*-rich diet (100 %), while mortality was not a toxicity bioindicator. It is concluded that the repelling ability and growth of *E. fetida* acted as early warning signals to predict the environmental risk of *Sargassum*. However, an aspect that deserves further research is the delayed effects of the prolonged exposure of *E. fetida* to soil with high *Sargassum* levels.

